# The *microRNA476a*/RFL module regulates adventitious root formation through a mitochondria-dependent pathway in *Populus*

**DOI:** 10.1101/852178

**Authors:** Changzheng Xu, Yuanxun Tao, Xiaokang Fu, Li Guo, Haitao Xing, Chaofeng Li, HuiLi Su, Xianqiang Wang, Jian Hu, Di Fan, Vincent L. Chiang, Keming Luo

## Abstract

Adventitious root (AR) formation at the base of stem cuttings determines the efficiency of clonal propagation for woody plants. Many endogenous and environmental factors influence AR formation. However, our knowledge about the regulation of AR development by mitochondrial metabolism in plants is very limited. Here we identified *Populus*-specific *miR476a* as a novel regulator of wound-induced adventitious rooting via orchestrating mitochondrial homeostasis in poplar. *MiR476a* exhibited inducible expression during AR formation and directly targets several *Restorer of Fertility like* (*RFL*) genes encoding mitochondrion-localized pentatricopeptide repeat proteins. Genetic modification of *miR476*-*RFL* expression revealed the *miR476*/*RFL*-mediated dynamic regulation of mitochondrial homeostasis on AR formation in transgenic poplar. Furthermore, mitochondrial perturbation via exogenous chemical inhibitor validated that the *miR476a*/*RFL*-directed AR formation depended on mitochondrial regulation though modulating the auxin pathway. Our results established a miRNA-directed mitochondrion-auxin signaling cascade required for AR development, providing novel insights into the understanding of mitochondrial regulation on plant developmental plasticity.

## Introduction

Adventitious roots (ARs) post-embryonically develop from non-root tissues in plants including hypocotyls, leaves, shoots, and reproductive organs (Bellini et al., 2014). Shoot-borne ARs occur normally in many monocotyledons, especially cereals such as rice and maize, which are essential for establishment of a fibrous root system in these plants (Hochholdinger et al., 2004; Bellini et al., 2014). In many plant species, AR emergence is usually induced by environmental changes such as flooding and dark-light transitions (Lorbiecke and Sauter, 1999; Sorin et al., 2005), or artificial stimuli, including wounding and hormone treatment (Ahkami et al., 2009; Sukumar et al., 2013). Cutting propagation through detached branches or stem is widely used in horticulture and forest nurseries (Davis and Becwar, 2007). As the first developmental event after cutting, adventitious rooting determines explant survival and efficiency of clonal propagation (Legué et al., 2014). AR formation is a wounding-triggered plant regeneration known as *de novo* root organogenesis that occurs when detached organs meet suitable conditions (Xu and Huang, 2014). AR formation starts with specification of root founder cells that undergo a few events, including cell division, patterning for AR primordium establishment and AR outgrowth (Legué et al., 2014). Genetic and molecular studies suggest that adventitious rooting requires coordinated and dynamic regulation of hormonal crosstalk and transcription factors (Bellini et al., 2014).

Auxin is considered as the master player and hub of signaling network during AR formation (Lakehal and Bellini, 2019). Indole-3-acetic acid (IAA) and its synthetic analogs are widely applied in AR induction for vegetative propagation (Preece, 2003). Auxin-mediated regulatory framework of wound-induced AR formation has been established in *Arabidopsis* involving auxin local synthesis, polar transport and signal transduction (Bellini et al., 2014). YUCCA1/4-mediated auxin biosynthesis promotes the fate transition of rooting competent cells in leaf explants of Arabidopsis (Chen et al., 2016). A wound-triggered signaling cascade mediated by jasmonate (JA) signaling was recently identified to stimulate auxin biogenesis for adventitious rooting (Zhang et al., 2019). JA-induced ETHYLENE RESPONSE FACTOR 109 directly activates the transcription of *ANTHRANILATE SYNTHASE α1* encoding a tryptophan biosynthetic enzyme for auxin production (Zhang et al., 2019). Increased auxin polar transport labeled by radioactive IAA was detected around the position of excision where ARs develop, in line with AR deficient phenotypes resulting from impaired polar auxin transport (Sukumar et al., 2013). Wound-induced auxin local maximum triggers the expression of *WUSCHEL RELATED HOMEOBOX* (*WOX*) *11* and *12* via the auxin responsive elements located within their promoters (Liu et al., 2014c). WOX11 and 12 act redundantly in specification of root founder cells, the first-step cell fate transition for AR emergence (Liu et al., 2014c). Auxin-directed activation of *WOX5/7* is required for cell patterning to establish AR primordia, the second step of cell fate transition during adventitious rooting (Hu and Xu, 2016).

In addition to Arabidopsis, the central role of auxin in AR formation was also well recognized in cereals (Orman-Ligeza et al., 2013; Bellini et al., 2014). OsWOX11, a key transcription factor (TF) of shoot-borne root emergence in rice, acts under the control of YUCCA-dependent auxin biosynthesis (Zhao et al., 2009; Zhang et al., 2018). *OsGNOM1* encoding a guanine-nucleotide exchange TF in rice regulates AR formation by modulating polar auxin transport (Liu et al., 2009). Gain-of-function of OsIAA23, a repressor of auxin signaling, blocks AR initiation (Ni et al., 2011). Some LOB and AP2 TFs were also identified as the key components in response to auxin during AR formation in rice (Inukai et al., 2005; Kitomi et al., 2011).

In woody species, *Populus FBL1* encoding a homologous auxin receptor of TIR was shown to regulate AR formation by interacting with Aux/IAA28 (Shu et al., 2019). Similar to *Arabidopsis* and rice, WOX5 and -11 also play crucial roles in the adventitious rooting process of *Populus* (Xu et al., 2015; Li et al., 2018). Moreover, the poplar AINTEGUMENTA LIKE1 (AIL1) and Response Regulator 13 (RR13) TFs were identified to mediate transcriptional programs during AR formation in *Populus* (Ramirez-Carvajal et al., 2009; Rigal et al., 2012). However, molecular understanding of AR morphogenesis in trees is still limited (Legué et al., 2014), despite importance for vegetative propagation, a currently most common practice for tree breeding and commercialization.

Regulation of plant growth and development, such as AR formation, often involves components other than hormones and TFs, such as microRNAs (miRNAs), for a collaborative and more specific mode of regulation. MiRNAs are a class of endogenous small RNAs that regulate an array of plant developmental programs, including adventitious rooting (Chen, 2009; D’Ario et al., 2017). Aberrant functions of some key components in miRNA biogenesis pathway, such as ARGONAUTE 1 in *Arabidopsis* and CROWN ROOT DEFECT 1 in rice, give rise to defective AR formation (Sorin et al., 2005; Zhu et al., 2019). AR-associated miRNAs and their targets are known to be part of auxin-directed molecular framework for adventitious rooting (Bellini et al., 2014). In *Arabidopsis*, *AUXIN RESPONSE FACTOR* (*ARF*) *6*/*8* and *17* antagonistically orchestrating AR formation are negatively regulated by *miR167* and *miR160*, respectively, by targeting their mRNAs for turnover (Gutierrez et al., 2009). Interactions of *miR167* and *miR160* with key TFs of the auxin signaling pathway are required for fine-tuning AR developmental plasticity (Gutierrez et al., 2009). Similarly, *miR160*-*ARF17* was characterized to regulate AR formation in a *Populus* hybrid, *P. deltoides* × *P. euramericana* (Liu et al., 2019). *MiR393* regulates AR formation in rice through repressing the expression of *TIR1*/*AFB2* auxin receptors (Bian et al., 2012). A *miR156*-mediated *SPL3*-*MADS50* module was recently identified to orchestrate AR formation in rice through affecting auxin signaling (Shao et al., 2019).

AR formation-associated miRNAs identified to date belong to the conserved miRNA families (Axtell and Bartel, 2005). There is a large number of nonconserved miRNAs in *P. trichocarpa*, of which functions are yet unknown (Lu et al., 2005). Of all the nonconserved miRNAs identified in *P. trichocarpa*, *miR476a* was found to have 20 predicted target genes, all encoding pentatricopeptide repeat (PPR) proteins (Lu et al., 2005). PPRs are typically targeted to mitochondria or chloroplasts and bind and influence organellar transcript expressions, leading to important effects on organelle biogenesis and ultimately plant development (Schmitz-Linneweber and Small, 2008; Barkan and Small, 2014).

Mitochondria are the energy producer in eukaryotic cells through coupling electron transport chain with oxidative phosphorylation for ATP generation (Millar et al., 2011). Apart from energy production and biosynthetic function, mitochondria are increasingly defined as a hub integrating signaling cascades for plant development and stress responses (Liberatore et al., 2016). The best example of mitochondrial signaling is cytoplasmic male sterility (CMS), a pathway leading to microsporogenesis defects and pollen abortion in many plant species (Carlsson et al., 2008). A more recent study revealed that fission, a dynamic behavior of mitochondria, modulates mitochondrial status in tapetal cells and thereby contributes to pollen viability, providing a direct link between mitochondrial homeostasis and pollen development (Chen et al., 2019). Mitochondrial dynamic changes affecting seed germination were also well documented (Law et al., 2012; Paszkiewicz et al., 2017). What specific roles mitochondrial homeostasis plays during root formation is unknown. The association of miRNAs with organellar PPRs allows us to begin to understand such roles, leading to new insights into a combined regulation of miRNAs, TFs of auxin signaling and mitochondrial homeostasis for AR formation and development.

Here, we identified that *miR476a*, a member of the *miR476* family, functions as a novel regulator of wound-induced AR formation via orchestrating mitochondrial homeostasis in *P. tomentosa*. *MiR476a* directly targets several mitochondrion-localized PPRs that belong to *Restorer of Fertility like* (*RFL*) genes. Genetic modification of *miR476a*-*RFL* expression and chemical perturbation of mitochondrial function revealed that the *miR476*/*RFL*-mediated dynamic regulation of mitochondrial homeostasis influenced AR formation via modulating the auxin pathway. Our results established a miRNA-directed mitochondrion-auxin cascade required for AR development in plants, especially in woody species.

## Results

### *MiR476a* positively regulates wound-induced AR formation in *Populus tomentosa*

The *miR476* family in the annotated *Populus trichocarpa* genome comprises two members that display high identity in precursor and mature sequences (*PtrmiR476a* and -*b*; Figure S1A). A pair of *miR476* homologs was also identified in *P. tomentosa*, and designated as *PtomiR476a* and -*b*. *PtomiR476*s exhibit high similarity in sequence and secondary structure to their *P. trichocarpa* counterparts (Figures S1A and S1B). We generated *PtomiR476a*-overexpressing poplar transgenic plants, and selected two lines (*miR476a*-OE-L1 and -L4) that had the highest transgene expression levels for further analyses (Figure S2A). Not only shoots but also a large number of roots were regenerated from the transgenic calli on shoot induction medium (SIM) (Figure S2B), suggesting that *miR476a* may enhance rooting capability, possibly even in the absence of rooting hormones. We then tested AR formation, the *de novo* root organogenesis, from wound-induced or cuttings of transgenic and wild type (WT) shoots and leaves in cultures without any hormones (Figures 1A to 1D). Shoot stem-derived ARs started to emerge 5 to 6 days after cutting (DAC) in both transgenic lines, but the emergence was delayed by ∼2 days in WT (Figures 1A and 1C). The analysis of the cross-sections of stem bases undergoing AR formation revealed that AR primordia (ARPs) were already initiated at 3 DAC from cambial zone of *miR476a*-OE plants while visible until 5 DAC in WT (Figure 1E). The *miR476a*-OE shoots developed 3 to 4 times more stem-derived ARs than did WT (Figures 1A and 1D), consistent with the initiation of more ARPs in *miR476a*-OE than in WT (Figure 1E). Similarly, faster and more ARs were derived from *miR476a*-OE leaves than from WT leaves (Figures 1B to 1D).

**Figure 1.**
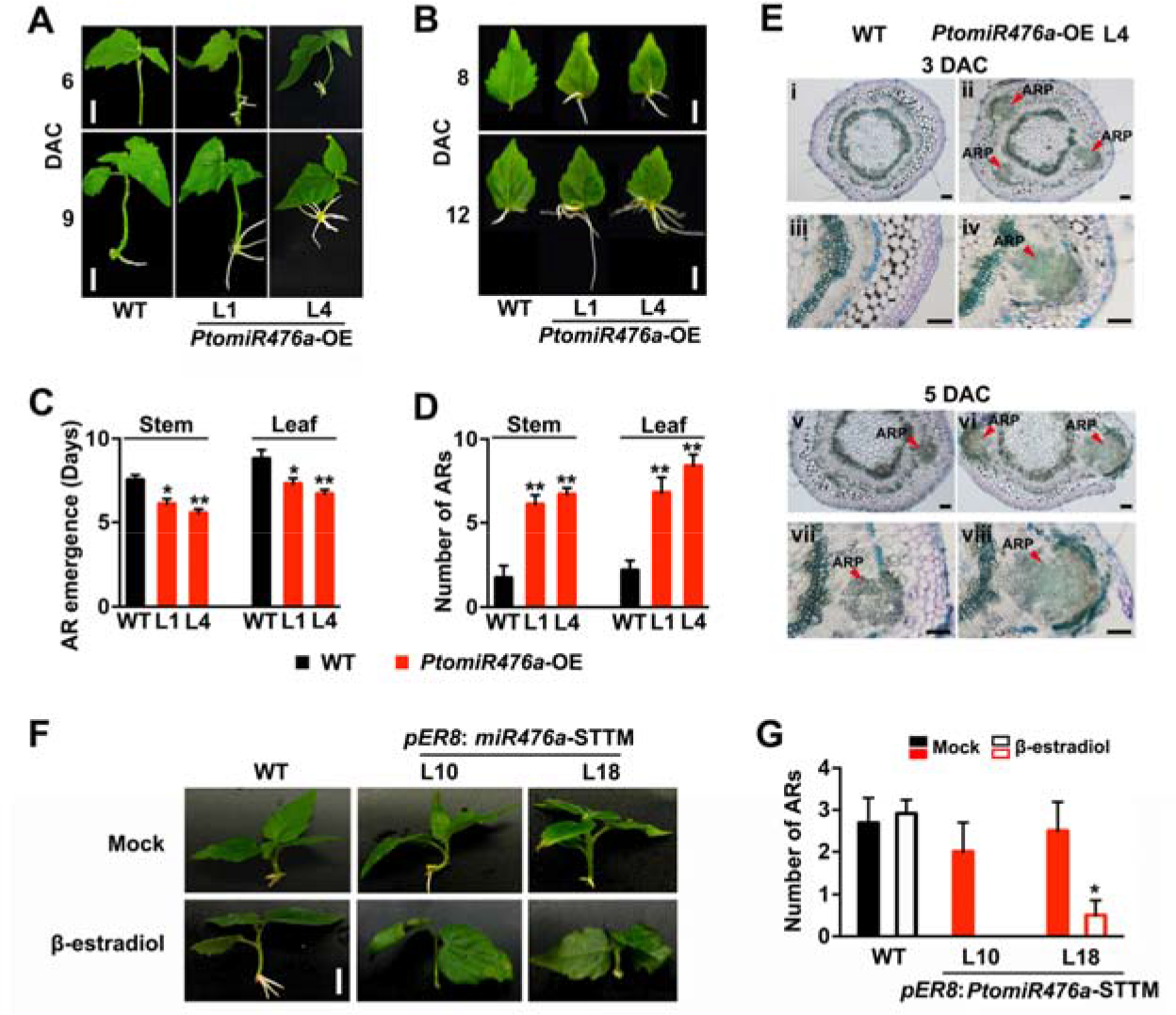
*MiR476a* positively regulates adventitious rooting capacity in poplar. A and B, phenotypes of ARs emerging from stem (A) or leaf (B) explants in wild-type (WT) and independent *miR476a*-OE poplar lines (L1 and 4). DAC: days after cutting. Bars = 1 cm (A) and 500 μm (B). C, quantification of the time for emergence of the first AR from stem and leaf explants. D, number of ARs derived from stem and leaf explants at 12 DAC. E, cross-sectioning and staining with toluidine blue of stem base at 3 and 5 DAC during AR formation in WT and the *miR476a*-OE line (L4). ARP: adventitious root primordium. Bars = 200 μm. F, phenotypes of AR formation in WT and independent *pER8*: *miR476a*-STTM transgenic lines (L10 and L18) under mock and β-estradiol treatment at 12 DAC. Bar = 1 cm. G, quantification of AR number at 12 DAC corresponding to F. Error bars represent SD. For C and D, asterisks indicate statistically significant differences in Student’s *t* test between WT and each transgenic line (*, *P* < 0.05; **, *P* < 0.01; n = 4). For F, asterisks indicate statistically significant differences in Student’s *t* test between mock and β-estradiol treatment for each line (*, *P* < 0.05; n = 3).

We also observed ectopic development of ARs in upper internodes of *miR476a*-OE plants cultivated either *in vitro* or in soil (Figure S2C), which do not occur in WT. The *miR476a*-OE shoot cuttings cultivated hydroponically also developed more ARs and from a more extended area than WT (Figure S2D). In potted soil, *miR476a*-OE plants developed a denser root system with enhanced lateral root branching but impaired root elongation compared with WT (Figures S2E, S2G and S2H). The total root biomass in *miR476a*-OE plants was significantly higher than that in WT (Figure S2F). Therefore, *miR476a* plays an important role in AR formation and some other key aspects of root organogenesis.

To further validate the *miR476a*-mediated regulation on AR formation, Short Tandem Target Mimic (STTM)-based knockdown (Bian et al., 2012) of *PtomiR476a* driven by the *CaMV 35S* promoter was performed (Figure S3A). However, the transgenic calli were difficult to induce roots on root induction medium (RIM) for generation of viable transgenic plants (Figure S3B), suggesting that constitutive knockdown of *miR476a* is lethal to poplar. Hence, we generated the *pER8*:*miR476a*-STTM transgenic plants that would confer the knockdown of *PtomiR476a* driven by a β-estradiol inducible promoter (Zuo et al., 2000) (Figures S3A and S3C). We selected two lines of *miR476a*-STTM transgenics (L10 and L18) with the strongest knockdown for testing *de novo* root organogenesis. Both lines displayed significantly retarded AR emergence when exposed to β-estradiol (Figure 1F). Compared with the mock, the transgenic L10 failed to form any ARs at 12 DAC, and L18 was 80% less efficient in generating ARs under the inducible knockdown of *miR476a* (Figure 1G). Both overexpression and knockdown experiments reveal that *miR476a* is required for AR formation in poplar.

### Dynamically inducible expression of *miR476a* during AR formation

To explain the *miR476a*-dependent AR phenotypes, tissue-specific and AR-associated expression patterns of *miR476a* were assayed in *P. tomentosa* (Figure 2). Quantitative PCR (qPCR) showed that *miR476a* was more abundantly expressed in roots, stems and petiole relative to leaves (Figure 2A). Similar expression patterns were detected by histologically staining with the GUS reporter driven by the *PtomiR476a* promoter (Figure 2B). In stems, *PtomiR476a* was exclusively expressed in cambial cells (Figure 2B), where ARs usually initiate in poplar (Rigal et al., 2012). High expression of *PtomiR476a* was also detected in both root tips (Figure 2Biv) and stele tissues (Figure 2Bv) of ARs. We next performed time-course analysis of the histological GUS activity during wound-induced AR formation (Figures 2C to 2E). The *PtomiR476a* expression remained low within 1 DAC at the base of stem cuttings, but drastically increased reaching a maximum at 6 DAC where the activity was ∼6 times of that at 1 DAC (Figures 2C and 2D). At 8 DAC, the activity decreased (by 62% compared to the maximum activity) along with the AR emergence (Figures 2C and 2D). GUS staining of cross-sections of stem cuttings validated the progressively increased expression of *miR476a* during AR formation (Figure 2E). The quantified promoter activity showed that *miR476a* appeared to be more abundantly accumulated at the basal area of emerged ARPs (Figure 2E). The inducible expression of *miR476a* during AR formation is consistent with the positive role for *miR476a* in regulating adventitious rooting capacity.

**Figure 2.**
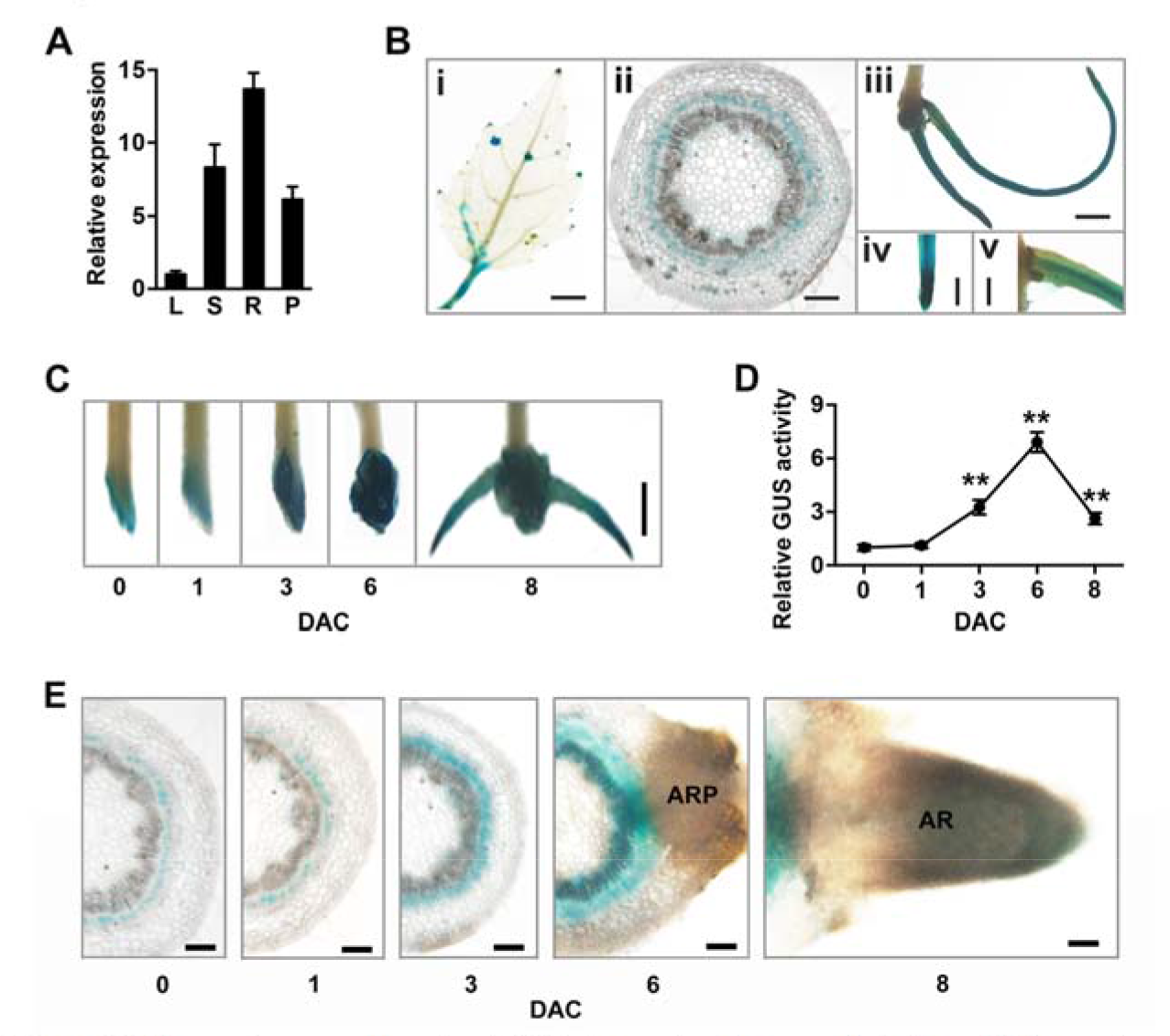
Expression patterns of *miR476a* in poplar tissues and during AR formation. A, tissue-specific expression of *miR476a* assayed by qRT-PCR in poplar. L: leaf blade; S: stem; R: root; P: petiole. B, histological staining of tissues of the transgenic poplar line harboring GUS reporter gene driven by the promoter of *miR476a*. i: leaf blade and petiole; ii: cross-sections of stem base; iii: adventitious roots; iv: adventitious root tip; v: upper part of adventitious roots close to stem. Bars = 5 mm (i and iii), 500 μm (ii), and 2 mm (iv and v). C, histological staining of stem bases close to cutting position of the GUS reporter line driven by the promoter of *miR476a* during AR formation. Bar = 5 mm. D, time-course quantitative assays of the GUS activity driven by the promoter of *miR476a* at stem bases during AR formation corresponding to C. E, histological staining of cross-sections of stem bases close to cutting position of the GUS reporter line driven by the *miR476a* promoter during AR formation. Bars = 500 μm. Error bars represent SD. For D, asterisks indicate statistically significant differences with respect to 0 DAC (Student’s *t* test; **, *P* < 0.01; n = 3).

### A group of RFL-PPR genes are targeted by miR476a during AR formation

Several *pentatricopeptide repeat* (*PPR*) genes were previously shown to be the potential targets of *miR476*s in *P. trichocarpa* (Lu et al., 2005). Based on the updated *P. trichocarpa* genome sequences (V3.0), we identified a total of 36 *PPR*s encoding Restorer of Fertility Like (RFL) proteins (Fujii et al., 2011) and designated according to their chromosomal locations (Figure S4). Using a similar prediction algorithm (Lu et al., 2005), 11 putative *miR476a* targets with high reliability (score ≤ 1.5) were identified from the 36 *RFL*s (Figures S4 and S5A). All 11 predicted *miR476a*-targeting *RFL* genes belong to the same phylogenetic subclade (Figure S4) and their counterparts were cloned from *P. tomentosa*. There are 3 or 4 unpaired bases between *PtomiR476a* and the predicted binding sites harbored by these *PtoRFL*s except for *PtoRFL27* (Figure S5B).

We next constructed degradome libraries from the WT and *miR476a*-OE lines for single-ended Illumina sequencing to provide evidence for the *miR476a*-targeted *RFL* cleavage (Figure S6A; Dataset S1). The degraded fragments, the cleavage of which occurred within the predicted binding sites of *miR476a*, were detected for 6 of the 11 predicted targets, i.e. *RFL1*, -*16*, -*17*, -*18*, -*26*, and -*34*, in both libraries (Figure S6A). These *RFL* transcripts contained 3 bases unpaired with *miR476a* within their predicted binding sites (Figure S5B). Due to the identical sequences of the predicted *miR476a* binding sites and flanking regions in *RFL1*, -*16*, -*18*, and -*34*, the fragments mapped to these *RFL*s could not be differentiated (Figure S6A). The cleaved fragments derived from these 6 *RFL*s accumulated in much higher frequencies in *miR476a*-OE plants than in WT (Figure S6A). Consistently, all of the 6 *RFL*s displayed significantly attenuated expression levels in *miR476a*-OE lines (Figure 3A). Other than these 6 *RFL*s, no accumulation of the cleaved fragments was detected for the other 5 predicted targets (*RFL19*, -*27*, -*30*, -*32*, and -*33*; Figure S6A). Moreover, the expression of these 5 *RFL*s was not modulated in *miR476a*-OE lines (Figure S6B). These results provide strong evidence that the 6 *PtoRFL*s, i.e. *PtoRFL1*, -*16*, -*17*, -*18*, -*26*, and -*34*, are most likely the authentic targets of *PtomiR476a*.

**Figure 3.**
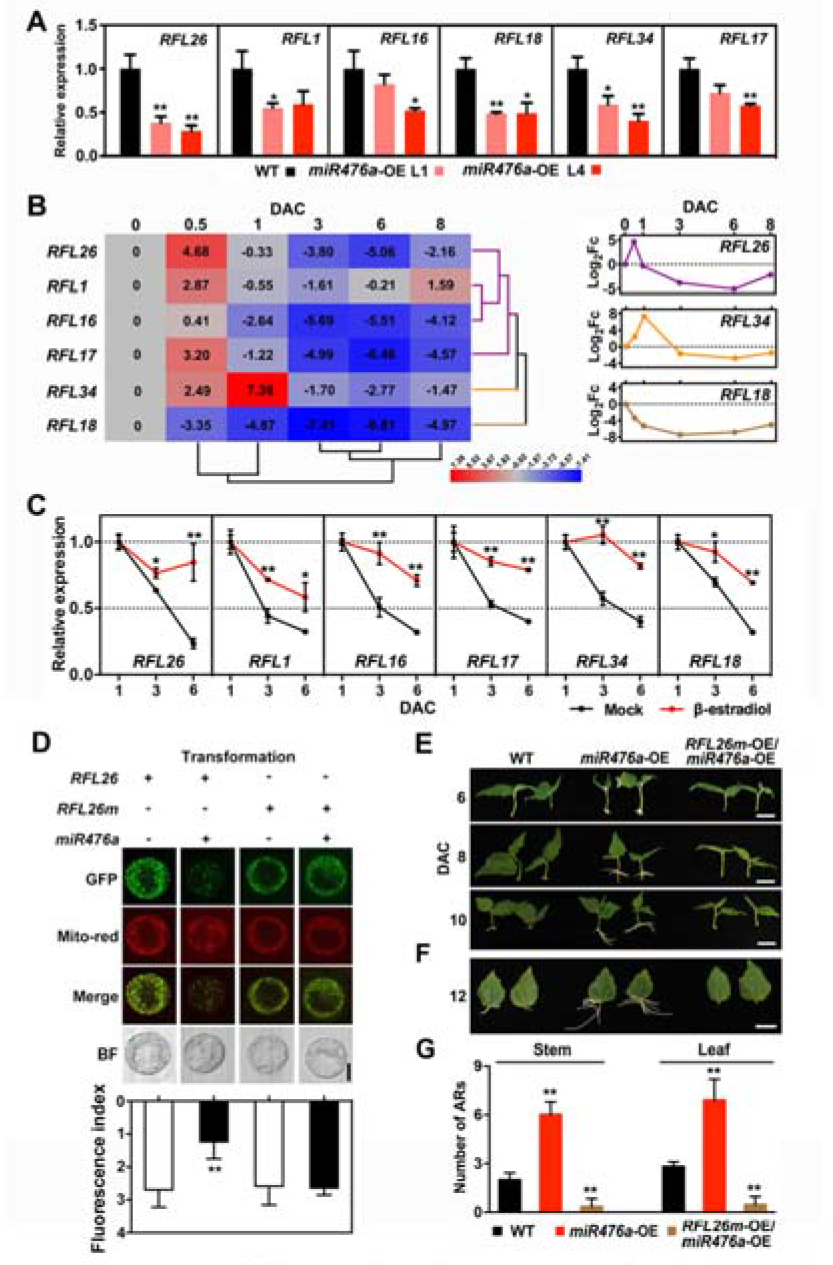
A group of *RFL* genes mediate *miR476a*-dependent AR formation in poplar. A, expression levels of *RFL1*, *16*, *17*, *18*, *26*, and *34* in WT and *miR476a*-OE lines determined by qRT-PCR. Stem bases of seedlings cultivated in vitro were collected for RNA extraction. *Ubiquitin* was used as a reference gene. The values of WT were normalized to 1. B, time-course expression assays of *miR476a*-targeted *RFL* genes during AR formation by qRT-PCR. For each gene, expression levels are normalized to that of 0 DAC and transformed to log2 values for heatmap analyses (leaf panel). Hierarchical clustering revealed three time-course expression patters of these *RFL* genes during AR formation, represented by those of *RFL26* (purple), -*34* (orange), and -*18* (brown) in the right panel. Fc: fold of change. C, expression of *miR476a*-targeted *RFL* genes in *pER8*: *miR476a*-STTM line under mock and β-estradiol treatment. D, experimental validation of the *miR476a*-targeted cleavage of *RFL26*. The plasmid of *miR476a*-OE was transiently transformed into the protoplasts isolated from roots of transgenic Arabidopsis harboring ectopically expressed *RFL26*-GFP and *RFL26m*-GFP. *RFL26m* represents the mutated *RFL26* gene harboring the impaired *miR476a*-binding site as indicated in Figure S10A. Red fluorescence was emitted from mitochondrial dye mito-tracker Red (Mito-red). The fluorescence was detected via confocal microscope (Olympus) and quantified by FV10-ASW software (Olympus). Bar = 10 μm. E and F, phenotypes of ARs derived from stem (E) and leaf explants (F) of overexpressing *RFL26m* in the *miR476*-OE line. Bars = 2 cm. G, quantification of AR number of overexpressing *RFL26m* in the *miR476*-OE line at 12 DAC. Error bars represent SD. For A and G, asterisks indicate statistically significant differences in Student’s *t* test between WT and each transgenic line (*, *P* < 0.05; **, *P* < 0.01; n = 4). For C, asterisks indicate statistically significant differences with respect to mock for each time point (Student’s *t* test; *, *P* < 0.05; **, *P* < 0.01; n = 5). For D, asterisks indicate statistically significant differences with respect to *RFL26* and *RFL26m* transformants, respectively (Student’s *t* test; *, *P* < 0.05; n = 5).

Given the inducible expression of *miR476a* during AR formation, transcript profiles of the *miR476a*-targeted *RFL*s were also determined during this process (Figure 3B). Clustering of the time-course assay datasets revealed various expression patterns of the *miR476a*-targeted *RFL*s during AR formation (Figure 3B). The first pattern shared by *RFL1*, -*16*, -*17*, and -*26* underwent dramatic increase of expression within 0.5 DAC and then rapid decrease during the following 1 - 5 DAC. *RFL34* exhibited the highest expression at 1 DAC. In contrast, the expression of *RFL18* was continuously decreased throughout this process. The decreased expression of the *miR476a*-targeted *RFL*s between 1 and 6 DAC coincided with the increased accumulation of *miR476a* (Figures 2C-E and 3C).

Since wounding is considered as a necessary stimulus for the initiation of ARs^48^. We speculated that both wounding signals and *miR476a* might regulate the stage-specific expression patterns of these *RFL*s. To test this idea, a time-course assay was performed in wounded leaves that were not detached for adventitious rooting. The results revealed that the expression of *RFL1*, -*16*, -*17*, -*26*, and -*34* was induced under wounding while that of *RFL18* was repressed (Figure S7A). The wounding responsiveness of these *RFL* genes mimicked the increase of their expression levels within 0.5-1 DAC during AR formation (Figures S7A and 3C). The qPCR assays in *pER8*:*miR476a*-STTM transgenic plants showed that the expression decrease of these *RFL* genes was significantly compromised when the knockdown of *miR476a* was triggered (Figure 3C). Therefore, the repressed expression of these *RFL* genes after 1 DAC largely depended on the *miR476a* expression during AR formation.

### RFL26 mediates *miR476a*-dependent AR formation in *P*. *tomentosa*

Pfam analyses^49^ showed that all the 6 validated *PtomiR476a* targets harbore 6 or 7 canonical PPR motifs (Figure S8), demonstrating that these RFL proteins belong to the P-type PPR subfamily^50^. Moreover, these RFL proteins displayed high probability of mitochondrial localization (Figure S8). Close phylogenetic relationship, conserved protein domains and common *miR476a*-directed transcript cleavage implied similar biological roles of these *RFL* genes. Therefore, we decided to focus on one target for a more thorough investigation. Since among the 6 validated targets, *RFL26* exhibited the highest frequency of cleavage by *miR476a* and the strongest transcript repression in *miR476a*-OE lines (Figures S6A and 3A), we selected it to further explore the *miR476a*-mediated genetic and molecular context of AR formation.

RNA *in situ* hybridization showed that the *RFL26* transcripts were preferentially accumulated in cambial zone at 1 DAC while in ARP tissues at 6 DAC (Figure S9). The *RFL26-GFP* fusion was stably introduced into *Arabidopsis*, and the GFP signals in the root cells were overlapped with those derived from the mitochondrion-specific fluorescent dye (Figure S10A), validating the mitochondrial localization of RFL26. The root protoplasts harboring RFL26-GFP were isolated and transiently transfected with *miR476a* (Figure 3D). Fluorescence detection revealed the significantly reduced abundance of RFL26 protein when *miR476a* was introduced into cells (Figure 3D). In contrast, RFL26 protein accumulation was not declined when the *miR476a*-binding site within the *RFL26* transcript was disrupted by mutagenesis (Figures 3D and S10B). As a control, RFL30, encoded by a *RFL* gene that could not be targeted by *miR476a* (Figure S6), did not exhibit the *miR476a*-inhibited accumulation (Figure S10C). These results suggested the *PtomiR476a*-directed post-transcriptional regulation of *PtoRFL26* encoding a mitochondrion-localized typical RFL-PPR protein.

To establish the genetic context of *miR476a*-RFL during AR formation, *RFL26m*, a *miR476a*-resistant *RFL26*, was introduced into the WT and *miR476a*-OE backgrounds (Figure S11A). Phenotypic analyses showed that *RFL26m* led to significantly delayed emergence and compromised number of ARs derived from stems and leaf explants, and even weaker rooting capacity than WT (Figures 3E to 3G and S11B). Functional complementation revealed the negative regulation of *miR476a*-targeted *RFL26* on AR formation.

### The *miR476a*-RFL module modulates functional homeostasis of mitochondria

*RFL* genes identified in rice, rape and *Petunia* encode the mitochondrion-localized PPR proteins that suppress the expression of mitochondria-encoded genes responsible for cytoplasmic male sterility (Bentolila et al., 2002; Kazama et al., 2008; Uyttewaal et al., 2008). A number of mitochondrial-encoded genes displayed differential expression levels in *miR476a*-OE and *RFL26m*-OE lines compared with WT (Figure S12), implying the functional modulation of mitochondria depending on the *miR476a*-RFL module. ATP content and NADH/NAD^+^ ratio were significantly increased in the *miR476a*-OE lines compared with WT, while decreased in *RFL26m*-OE lines (Figures 4A and 4B). To further uncover mitochondrial homeostasis mediated by *miR476a*/*RFL*, time-course measurement of ATP content was conducted during AR formation (Figures 4C and 4D). A reduction of ATP content within 0.5 or 1 DAC followed by a dramatic increase along with ARP emergence at 6 DAC was detected (Figure 4C). However, the reduction of ATP level was absent at the early stage of AR formation due to the overexpression of *miR476a*, and higher ATP content was detected during the whole period of AR formation in the *miR476a*-OE plants than WT (Figure 4C). In contrast, a decrease in ATP accumulation was present during AR formation in the *RFL26m*-OE plants (Figure 4C). Similar changes were detected for NADH/NAD^+^ ratio among *miR476a*-*RFL26m* transgenic lines (Figure 4D). In addition, DAB staining and quantitative measurement revealed that H_2_O_2_, a molecule of reactive oxygen species (ROS), displayed preferential accumulation in *RFL26m*-OE lines, while compromised in *miR476a*-OE plants (Figures 4E and 4F). Taken together, these results revealed the antagonistically modulated mitochondrial functions by *miR476a* and its target *RFL*26 in *P. tomentosa*.

**Figure 4.**
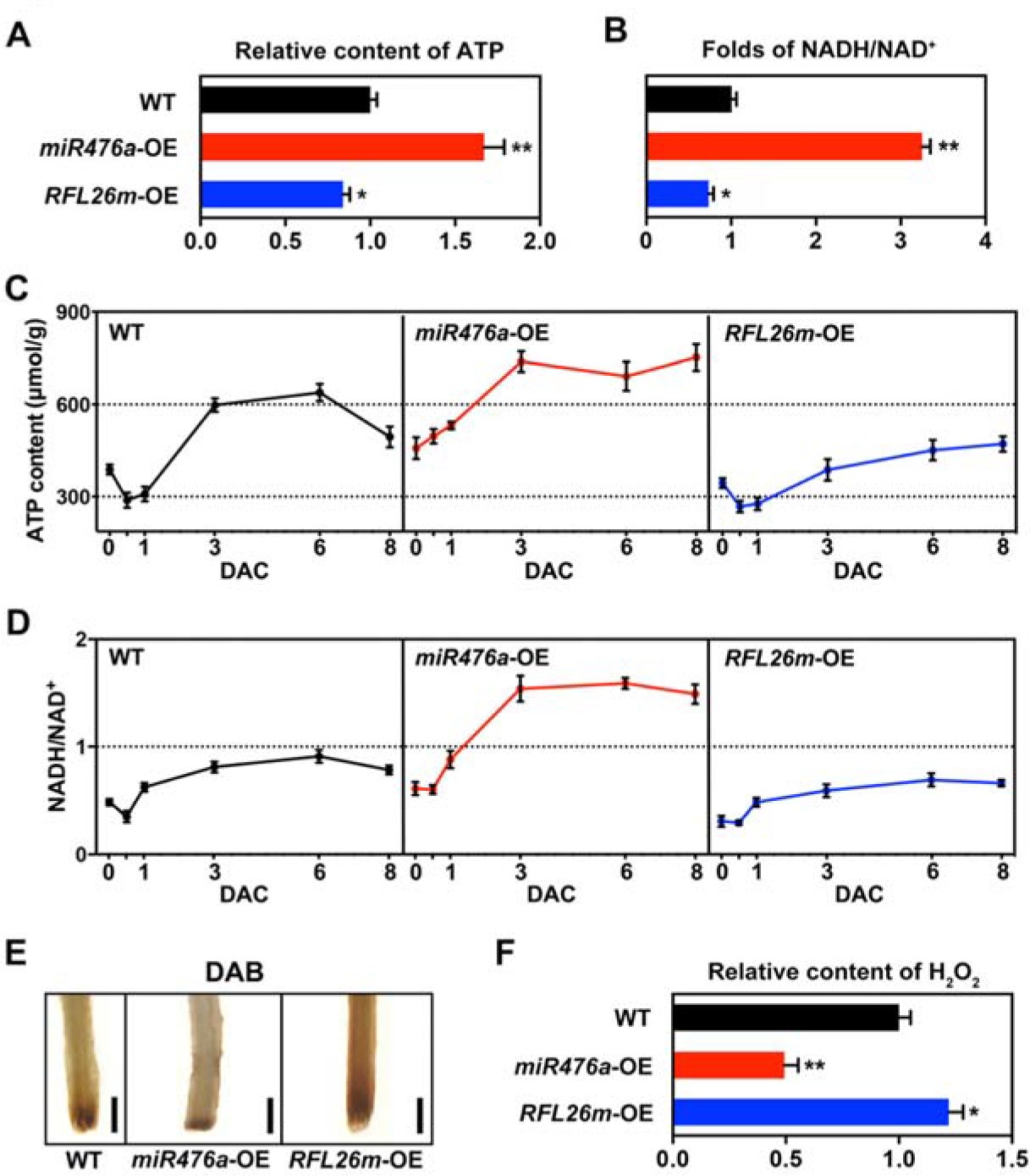
Mitochondrial homeostasis and dynamics modulated by *miR476a/RFL*. A and B, Quantitative assays of ATP (A) and NADH/NAD+ (B) in WT, miR476a-OE, and RFL26m-OE lines. Stem bases at 3 DAC were collected for measurements. C and D, Time-course quantification of ATP production (C) and NADH/NAD+ ratio (D) during AR formation in poplar. E, DAB staining of 3-DAC stem bases of WT, miR476a-OE, and RFL26m-OE lines. Scale bars = 0.5 cm. F, quantification of H_2_O_2_ content of 3-DAC stem bases of WT, miR476a-OE, and RFL26m-OE lines. Error bars represent SD. For A, B, and F, the values of WT are normalized to 1, and asterisks indicate statistically significant differences with respect to WT (Student’s *t* test; *, *P* < 0.05; **, *P* < 0.01; n = 3).

### AR formation requires *miR476a*/RFL-mediated mitochondrial regulation

Given the dynamic expression and coordinated regulation on mitochondria during AR formation, we questioned whether *miR476a*-driven mitochondrial modulation is required for adventitious rooting capacity. To test this hypothesis, exogenous application of rotenone, a chemical inhibitor of activity of mitochondrial respiratory chain complex (MRCC) I, was performed to determine the effect of mitochondrial respiration on AR development. ATP content was strongly inhibited and maintained at low levels in both WT and *miR476a*-OE lines after exposure to rotenone at 1 DAC during AR formation (Figure 5A). A similar reduction was detected for NADH/NAD^+^ in response to rotenone treatment (Figure 5B). AR development was suppressed by rotenone from shoot explants of both WT and *miR476a*-OE lines (Figure 5C). Accordingly, 2-3 days were delayed for AR emergence from WT and *miR476a*-OE explants under rotenone treatment (Figure 5D). The decreases of AR number were also detected for both genotypes (73.8% for WT and 77.3% for *miR476a*-OE; Figure 5E). These results indicated that functional homeostasis of mitochondria compromised by exogenous chemical inhibitor retarded AR formation, mimicking the *RFL26*-resulting AR defects. Therefore, mitochondrial regulation is required for adventitious rooting capacity in poplar.

**Figure 5.**
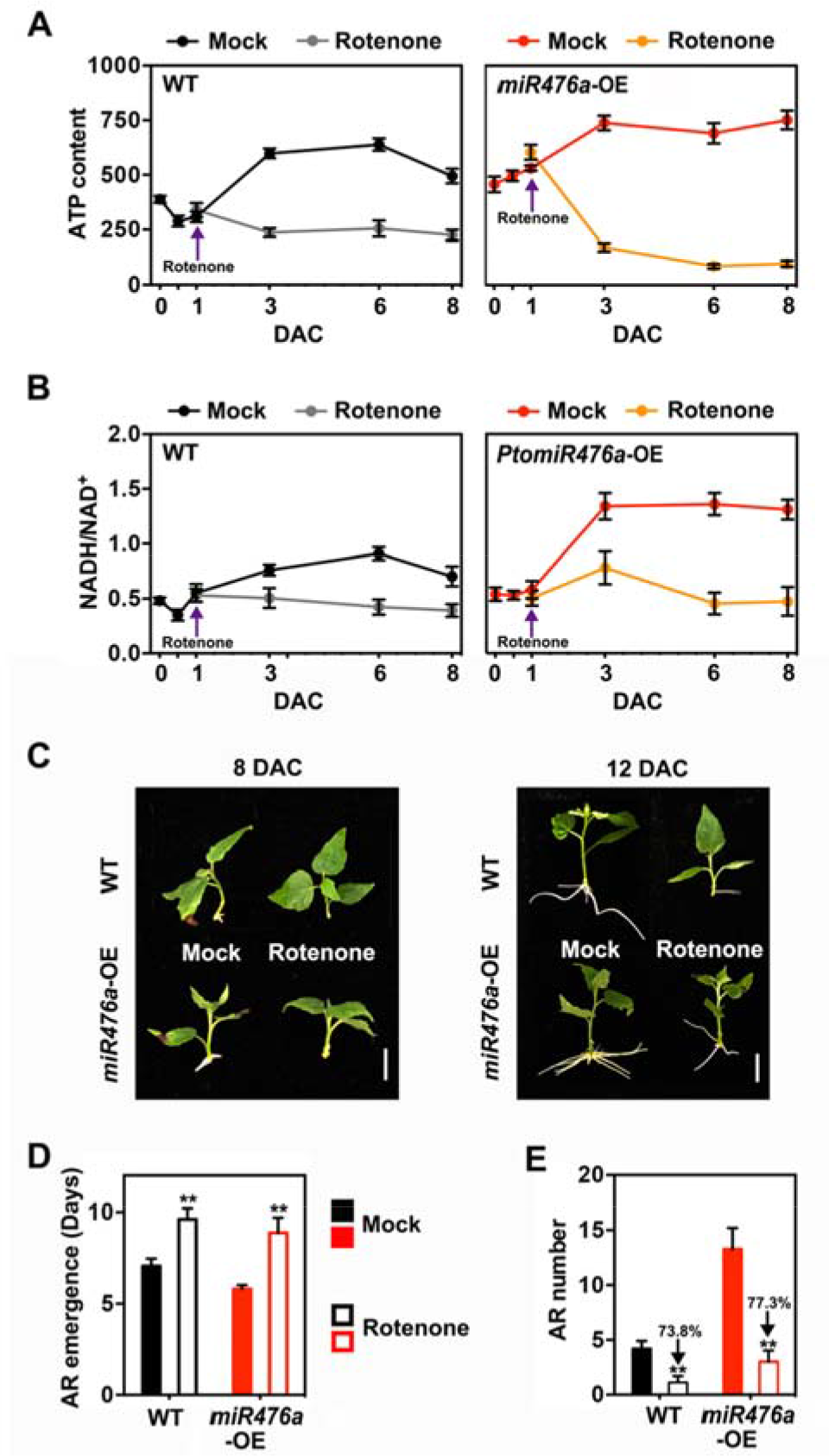
*MiR476a*-mediated AR formation depends on mitochondrial homeostasis. A and B, rotenone-inhibited ATP production (A) and NADH/NAD^+^ (B) during AR formation in WT and *miR476a*-OE lines. Rotenone (50 μM) was added for treatment at 1 DAC. C, AR phenotypes of WT and *miR476a*-OE lines under rotenone treatment at 8 and 12 DAC. Bars = 2 cm. D and E, quantification of the time for emergence of the first AR (D) and AR number (E) under rotenone treatment at 12 DAC in WT and *miR476a*-OE lines. Error bars represent SD. For D and E, asterisks indicate statistically significant differences with respect to mock (Student’s *t* test; **, *P* < 0.01; n = 5).

To examine whether mitochondrial regulation on adventitious rooting is conserved across woody and herbaceous plants, ATP quantification and mitochondrial perturbation were conduced using *Arabidopsis* leaf explants for AR formation. Similar to poplar, the *Arabidopsis* explants displayed 2-fold increase of ATP content at 4 DAC during AR formation (Figure S10A). The rotenone treatment resulted in dramatically retarded the emergence of ARs from leaf explants in *Arabidopsis* (Figures S10B and S10C). These results suggested conserved regulation of mitochondrial homeostasis on adventitious rooting in herbaceous species.

### *MiR476a*-mediated AR formation depends on auxin transport and homeostasis

To explore the *miR476a*-mediated molecular context of AR formation, comparative transcriptomic profiling was conducted *via* RNAseq, showing 484 differential expression genes (DGE) in *miR476a*-OE plants and WT (fold change ≥ 2, false discovery rate ≤ 1%; Figure S14A). Among them, 30 genes from different transcription factor families were identified to be regulated by *miR476a* overexpression (Figure S14B). These genes including *WUSCHEL-RELATED HOMEOBOX* (*WOX*) *4/5/11*, *AINTEGUMENTA LIKE* (*AIL*) *1*, and D-type *Cyclin* (*CYCD*) *3.1*, which are known to positively regulate AR formation in poplar (Karlberg et al., 2011; Rigal et al., 2012; Liu et al., 2014a), were preferentially expressed in *miR476a*-OE plants (Figure S14C). In contrast, these genes were down-regulated in both β-estradiol-induced *miR476a*-knockdown and *RFL26m*-OE lines (Figures S14D and S14E), indicating that they function downstream of the *miR76a*/RFL module during AR formation in poplar. The time-course assay of stem-derived AR formation revealed the increased transcript accumulation of *WOX5a* until the emergence of ARPs at 6 DAC in WT (Figure S14F), while its expression at high level in the *miR476a*-OE plants during the whole process of AR formation but at low level in the *RFL26m*-OE plants (Figure S14F. In addition, exogenous rotenone treatment could significantly suppress the expression of the *WOX* genes, validating mitochondrial regulation on *WOX*-mediated AR formation (Figure S14G).

A previous study in *P. tomentosa* showed that *PIN2* encodes a plasma membrane-localized efflux carrier essential for auxin polar transport, while *PIN5*s encode endoplasmic reticulum (ER) membrane-localized transporters affecting auxin cellular homeostasis (Liu et al., 2014b). The expression of *PIN2* and -*5b* were determined in *miR476a*- and *RFL26m*-OE lines (Figures 6A and 6B). Compared to WT, both *PIN* genes were preferentially expressed in *miR476a*-OE plants while weakly in *RFL26m*-OE plants (Figure 6A). Correspondingly, the expression levels of both *PIN*s were attenuated by the β-estradiol-induced *miR476a* knockdown (Figure 6B). The modulated expression of *PIN2* and -*5b* raised the likelihood that auxin homeostasis and polar transport function downstream of *miR476a*/RFL during AR formation. Compromising auxin homeostasis via application of 1-N-naphthylphthalamic acid (NPA), an inhibitor of auxin polar transport, was conducted to test this hypothesis. We found that no ARs emerged at 12 DAC under NPA treatment for both WT and *miR476a*-OE explants (Figure 6C). These results suggested that auxin polar transport is required for the *miR476a*/RFL-directed mitochondrial regulation of AR formation.

**Figure 6.**
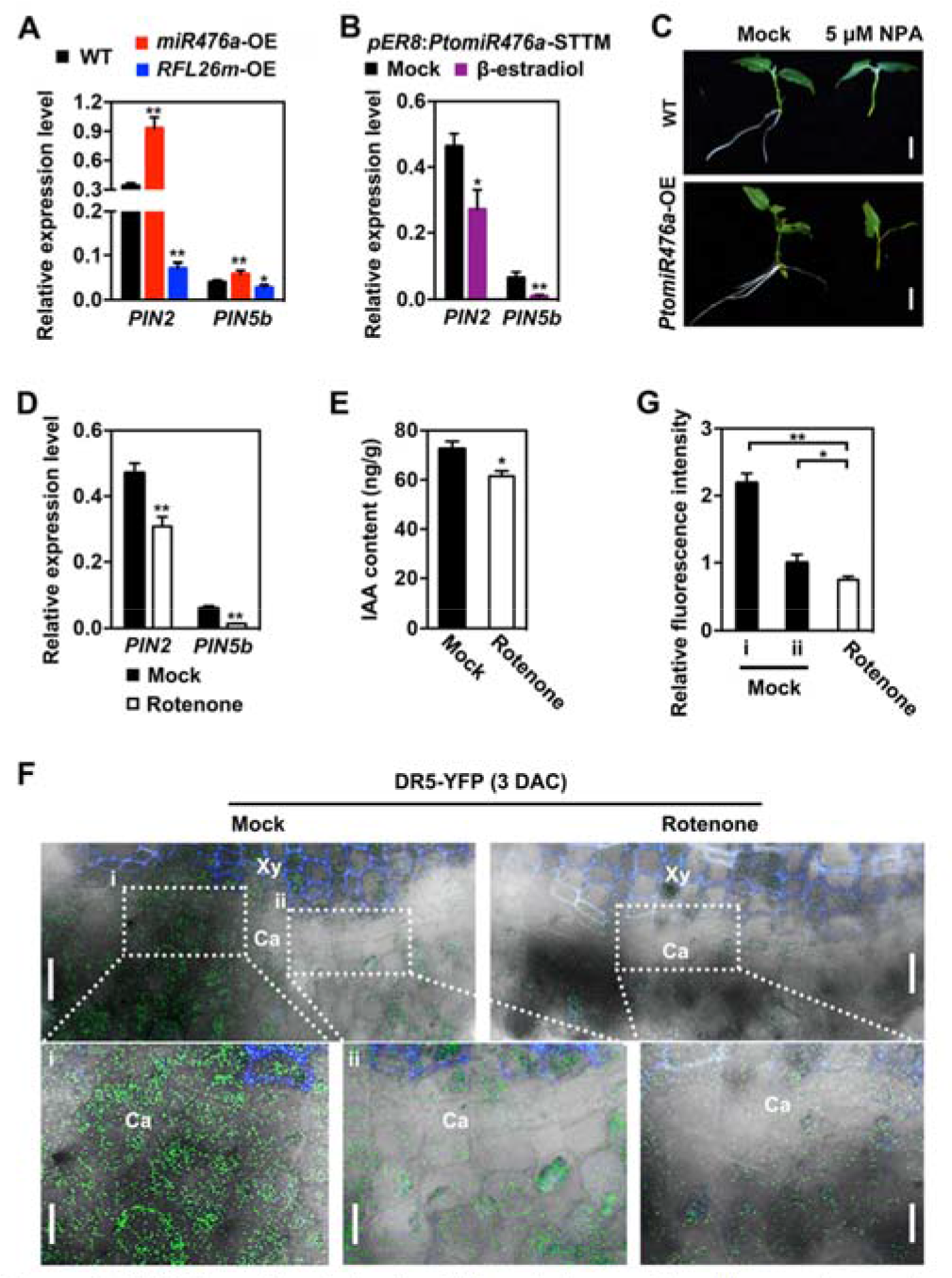
*MiR476a*-mediated mitochondrial regulation on adventitious rooting requires auxin pathway. A, expression levels of *PIN2* and *5b* in *miR476a*-OE and *RFL26m*-OE lines at 3-DAC stem bases assayed by qRT-PCR. B, expression levels of *PIN2* and *5b* in the *pER8*: *miR476a*-STTM line under mock and β-estradiol treatment. C, AR phenotypes of WT and *miR476a*-OE plants at 8 DAC under mock and 5 μM 1-N-naphthylphthalamic acid (NPA) treatment. D, expression levels of *PIN2* and *5b* under 50μM rotenone treatment for 3 days during AR formation. E, quantitative assays of indole-3-acetic acid (IAA) in stem bases under 50 μM rotenone treatment for 3 days during AR formation. F, fluorescent imaging of cross-sections of the *DR5*-*YFP* reporter line at 3 DAC under 50μM rotenone treatment. Red dashed circle indicates an area where YFP fluorescence is highly accumulated. Pf: phloem fibers; Ca: cambium; Xy: xylem. Bars = 20 μm. G, quantification of YFP fluorescence corresponding to H. The values of mock (i) are normalized to 1. Error bars represent SD. For A, asterisks indicate statistically significant differences with respect to WT (Student’s *t* test; *, *P* < 0.05; **, *P* < 0.01; n = 5). For B, D, E, and G, asterisks indicate statistically significant differences with respect to mock (Student’s *t* test; *, *P* < 0.05; **, *P* < 0.01; n = 3).

In consistence with the *miR476a*/RFL26-mediated regulation on *PIN2/5b*, the expression of both *PIN*s was suppressed by exogenous treatment of rotenone during AR formation (Figure 6D). This implied probable mitochondrial modulation of auxin transport and homeostasis, and IAA content was thereby examined in shoot explants for rooting under rotenone treatment (Figure 6E). Auxin content at stem bases of the explants on 3 DAC was reduced by 15.3% under rotenone treatment (Figure 6E), indicating negative effects of mitochondrial perturbation on auxin accumulation for rooting. Finally, the *DR5:YFP* reporter, a biosensor of in vivo auxin signaling, was introduced into poplar to investigate mitochondrial regulation on auxin pathway during AR formation (Figure 6F). Fluorescence detection of stem bases of shoot explants for AR formation revealed a preferential accumulation of auxin signaling at some position of cambial and neighboring zone at 3 DAC (Figure 6F). In contrast, the uneven distribution of auxin signaling at cambium disappeared in response to rotenone treatment (Figure 6F), and the fluorescence quantification showed the significantly rotenone-suppressed auxin signaling in comparison with the mock treatment (Figure 6G). Therefore, functional perturbation of mitochondria restricts auxin pathway during AR formation.

## Discussion

### MiRNA-directed dynamic posttranscriptional control of *RFL* genes synchronizes mitochondrial status with adventitious rooting

Mitochondria are the major organelle for primary metabolism and energy production in plant non-photosynthetic organs (Millar et al., 2011). A variety of developmental defects were identified in some mutants of genes encoding mitochondrion-localized proteins (Millar et al., 2011). The emerging role of mitochondrial homeostasis and dynamics has been characterized in a few developmental programs of *Arabidopsis* including seed germination, leaf senescence, and pollen development (Chrobok et al., 2016; Paszkiewicz et al., 2017; Chen et al., 2019). The loss-of-function of the mitochondrial pyruvate dehydrogenase E1 subunits led to root morphogenesis defects (Ohbayashi et al., 2019), suggesting the regulation of metabolic adjustment in mitochondria on root development. However, molecular connection and action mode enabling coordination of *de novo* root organogenesis and mitochondrial homeostasis has not been established yet. Here we identified a miRNA-directed signaling pathway that coordinates mitochondrial functional balance with adventitious rooting capacity in *Populus*. *MiR476* was initially cloned in tension/compression-stressed woody tissues in *P. trichocarpa* (Lu et al., 2005). We found the induced enrichment of *PtomiR476a* in cambium and neighboring parenchyma cells where stem-derived ARs developed (Figure 2). This dynamic expression pattern was consistent with the positive regulation of *miR476a* on adventitious rooting capacity addressed by genetic evidence (Figure 1). Noticeably, the induction of *miR476a* was absent at the early stage after wounding (before 1 DAC; Figure 2). Therefore, *miR476a* functions as a stage-specific regulator of AR formation.

Lu et al. (2005) showed *miR476a*-directed cleavage of a *PPR* gene via 5’ RACE. We provided multiple lines of evidence that *miR476a* targeted a group of *RFL* genes for transcript turnover to allow suppression of their expression (Figures 3 and S6). In addition to genetic complementation (Figures 3D to 3F), spatiotemporal patterns of *miR476a* and its *RFL* targets addressed their antagonistic regulation on adventitious rooting (Figures 2 and 3). Functional status of mitochondria characterized by energy production was activated by *miR476a* whereas suppressed by *RFL*s (Figures 4A and 4B). We found a stage-specific modulation of mitochondrial homeostasis during wound-induced AR formation in poplar (Figures 4D and 4E). The wound-induced expression of *RFL*s accounts for the decreased mitochondrial functions within 1 DAC while the *miR476a*-attentuated accumulation of the *RFL* transcripts contributes to the increase of mitochondrial functions thereafter (Figure 4). *MiR476a*/*RFL*-mediated mitochondrial modulation allows us to divide the wound-induced AR formation into two sequential phases, including wound response and root induction. Our results revealed that mitochondrial functional homeostasis is dynamically coordinated with wound-induced AR formation by the *miR476a*-directed posttranscriptional control of *RFL*s encoding mitochondria-localized proteins in poplar.

### Mitochondrial regulation of energy homeostasis is crucial for adventitious rooting

Due to the sessile lifestyle of plants, mitochondrial functionality for energy production requires dynamic flexibility depending on variable cellular functions (Schwarzlander and Finkemeier, 2013). For instance, mitochondria are immediately reactivated upon imbibition to support germination program of seeds for organogenesis of a new seedling (Paszkiewicz et al., 2017). Higher respiratory rate of mitochondria has been detected for root meristem than the other root tissues (Kuroiwa et al., 1992; Fujie et al., 1993). AR formation is a process of cell fate reprogramming for root organogenesis in parallel with active cell proliferation and differentiation (Legué et al., 2014; Pizarro and Díaz-Sala, 2018). A demand for high energy may exist to support these cellular behaviors, hinted by dynamic carbohydrate allocation at the base of cuttings during AR formation (Ahkami et al., 2009). Our results indicated that AR induction requires the elevated energy supply of mitochondria by the *miR476a*-restricted *RFL* expression (Figure 4). The genetic evidence was validated by pharmacological inhibition of mitochondrial functionality, which mimicked the *RFL*-resulting perturbation of energy production, and arrested *miR476a*-directed AR formation (Figure 5). The evidence of genetically or chemically modulated mitochondrial functionality suggested the key role of mitochondria-directed energy homeostasis in AR formation.

Similar to poplar, the elevated mitochondria-producing energy was detected for AR formation in *Arabidopsis*, the inhibition of which also impaired adventitious rooting capacity (Figure S13). These results revealed the conserved mitochondrial regulation on AR formation across herbaceous and woody plant species. However, *miR476* was considered as a *Populus*-specific miRNA family, due to its absence in the genome of *Arabidopsis* (Lu et al., 2005). It is thereby hypothesized that, although the conserved demand for mitochondria-generating energy, a variable action mode may function for coordinating mitochondrial homeostasis with adventitious rooting in Arabidopsis.

### Mitochondrial actions on AR formation requires retrograde regulation on auxin transport and homeostasis

Nucleus-organelle communication require bi-directional signals called anterograde and retrograde (de Souza et al., 2017). Retrograde signals confer feedback regulation from organelles including mitochondria on transcriptional responses in nucleus(Wagner et al., 2018). In addition to the miRNA-directed mitochondrial homeostasis, our results also showed that retrograde regulation on the auxin pathway delivers physiological alternations of mitochondria to direct AR formation (Figure 6). The *Arabidopsis mas* (*More Axillary Shoots*) mutant lacking the mitochondrial FTSH4 AAA-protease exhibits the compromised auxin accumulation or responsiveness, supporting a mechanistic link between mitochondrial functions and auxin (Zhang et al., 2014). Independently, using a chemical screening approach, two mitochondrion perturbing compounds sharing a common 3-(2-furyl) acrylate substructure were identified to inhibit auxin signaling(Kerchev et al., 2014). We found the modulated expression of *PIN2/5* encoding auxin transporters at the bases of cuttings in response to genetically or chemically orchestrated mitochondrial functionality (Figure 6). Due to exclusively inducible expression during AR formation, *PIN2/5* genes were considered to regulate polar transport and homeostasis of auxin for AR formation (Liu et al., 2014b). Importantly, exogenous treatment of auxin transport inhibitor and detection of auxin content and signaling under chemical perturbation of mitochondria validated the requirement of the retrograde regulation on auxin pathway (Figure 6).

Despite mitochondrial regulation on auxin homeostasis, the substances functioning as retrograde signals during AR formation are still unknown. ROS production in mitochondria is considered as a candidate signal of mitochondrial energy signaling (Schwarzlander and Finkemeier, 2013; Wagner et al., 2018). In the case of the *ftsh4* mutant, H_2_O_2_, a species of ROS, was identified to mediate mitochondrial regulation on auxin catabolism (Zhang et al., 2014). We also found *miR476a*-suppressing but *RFL*-promoting accumulation of H_2_O_2_ along with energy modulation during AR formation (Figure 4C). However, more lines of evidence are still required to elucidate the role of ROS in mitochondrial retrograde regulation during adventitious rooting.

Collectively, we proposed a model of the *miR476a*/RFL-directed mitochondrial regulation on adventitious rooting capacity in *Populus* that is defined as two sequential and interconnected stages (Figure 7). Wound at the bases of cutting explants induces the expression of several *RFL* genes encoding mitochondria-localized PPR proteins, and thereby restricts mitochondrial energy production that is unfavorable to AR initiation. Subsequently, the enrichment of *miR476a* defines the secondary phase for rooting induction. *MiR476a* directs the cleavage of the *RFL* transcripts for degradation, thereby suppressing the amount of the mitochondria-targeted RFL proteins. The *miR476a*-dependent suppression of RFLs causes the augmented mitochondrial status for energy supply indicated by increased ATP and NADH/NAD^+^ production whereas decreased ROS generation. The retrograde signals from mitochondria activate the nuclear expression of *PIN2*/*5b* encoding auxin transporters. PIN2/5b-modulated polar transport and homeostasis contribute to auxin maxima at rooting competent cells, and thereby drive AR formation via promoting the expression of AR-associated key genes. Our findings provide new insights into the mechanism by which a miRNA-directed signaling pathway dynamically coordinates mitochondrial functionality with adventitious rooting capacity in *Populus*.

**Figure 7.**
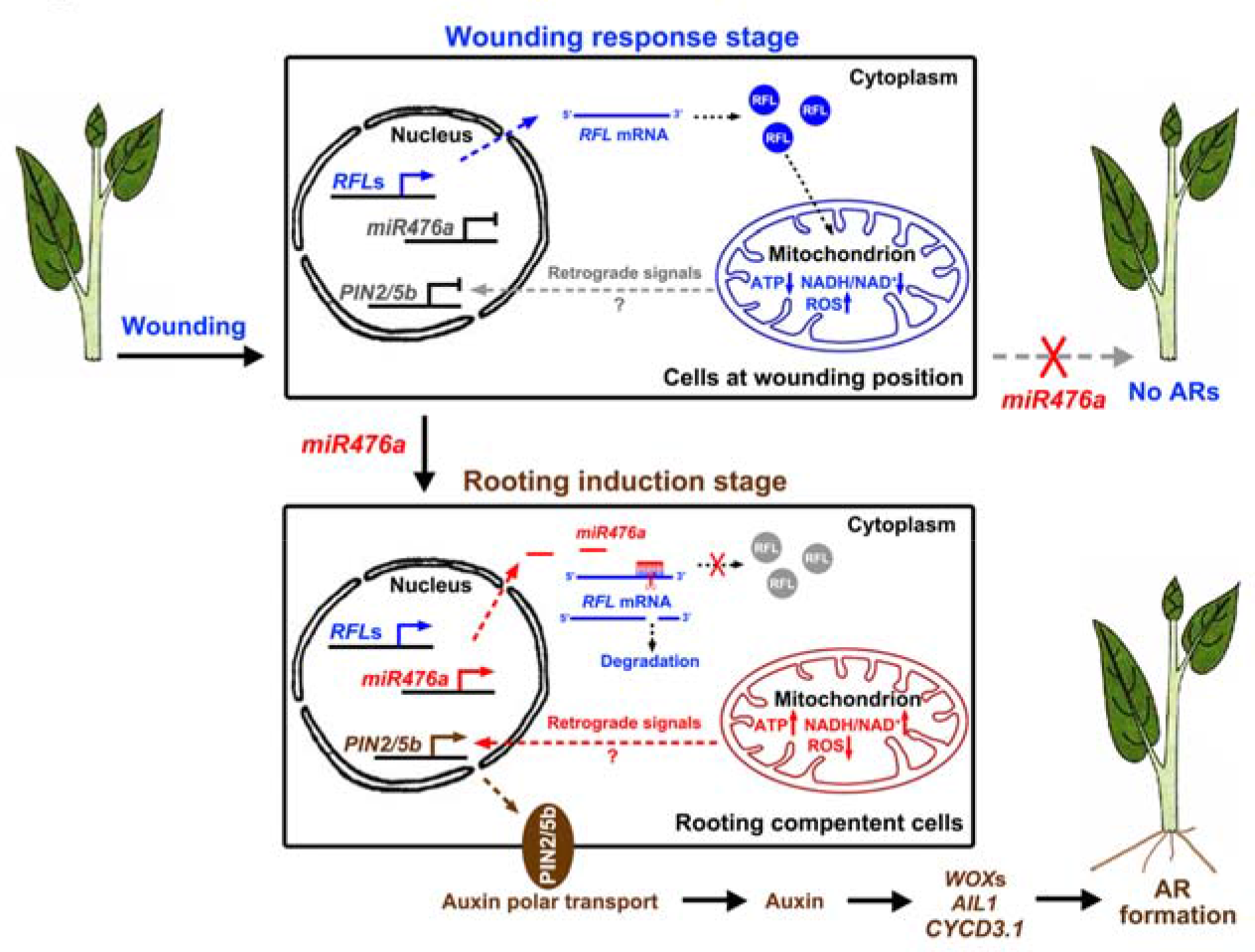
A proposed model for *miR476a*/*RFL*-dependent AR formation in *Populus*. The *miR476a*/*RFL*s-mediated mitochondrial functional modulation allows to divide the process of wound-induced AR formation into two sequential phases, including wounding response and root induction. At the first stage, wound signals at the bases of cutting explants induce the expression of several *RFL* genes encoding mitochondria-localized PPR proteins, and thereby restrict mitochondrial energy production that is unfavorable to AR initiation. At the second stage, the induction of *miR476a* results in the cleavage of the *RFL* transcripts for degradation, thus suppressing the amount of RFL proteins targeted into mitochondria. *MiR476a*-dependent suppression of *RFL*s causes the augmented mitochondrial status for energy supply indicated by increased ATP and NADH/NAD^+^ production whereas decreased ROS generation. The mitochondrial retrograde signals activate the expression of *PIN2*/*5b* auxin transporters, which modulated polar transport and homeostasis of auxin at rooting competent cells, and promote AR formation through elevating the expression of AR-associated key genes.

## Materials and methods

### Gene cloning and plasmid construction

The precursors of *PtomiR476a* and -*b* were amplified from genomic DNA of *P*. *tomentosa* using the primer pairs *Pre*-*PtomiR476a*-OE-*Bgl*II-fw/*Spe*I-rv and *Pre*-*PtomiR476b*-fw/rv (Table S1), respectively, and ligated into the pMD19 vector via TA-cloning for sequencing. The *miR476a* precursor was then subcloned into the pCAMBIA1302 vector driven by the *CaMV 35S* promoter via *Bgl*II/*Spe*I restriction sites. The STTM-based knockdown plasmid of *miR476a* was constructed as previously described (Bian et al., 2012; He et al., 2018). An oligonucleotide fragment for STTM containing the sequences of the *miR476a*-binding site with an insertion of AGA and an 88-bp spacer sequence was de novo synthesized (Figure S3A), and then inserted into the pCXSN vector under the control of the *CaMV 35S* promoter via *Bam*HI restriction sites (Chen et al., 2009). For inducible knockdown, the STTM fragment was amplified using the primers *PtomiR476a*-STTM-*Xho*I-fw/rv (Table S3), and ligated into the pER8 vector encoding a β-estradiol-inducible expression system (Zuo et al., 2000) via *Xho*I restriction sites. A 2973-bp promoter sequence upstream of the *miR476a* precursor was amplified from genomic DNA of *P. tomentosa* via the primer pair *Pro-PtomiR476a*-fw/rv (Table S3), and constructed into *Xcm*I-digested pCXGUS-P via TA-cloning to drive the expression of GUS (Chen et al., 2009). The full-length coding sequences of *RFL1*, *-16*, *-17*, *-18*, *-19*, *-26*, *-27, -30*, *-32*, *-33,* and *-34* were amplified from the *P. tomentosa* cDNA using primer pairs *PtoRFL*-CDS-fw/rv (Table S1), respectively, and constructed into pCXSN for sequencing via TA-cloning. For GFP tagging, the coding sequences of *PtoRFL26* and *PtoRFL30* without the stop codon were subcloned into pCAMBIA1302 with the hygromycin-resistant gene using the primer pairs CDS-*Nco*I-fw/noTAA-*Nco*I-rv for *PtoRFL26* and *30* (Table S1), respectively. For overexpression, the coding sequence of *PtoRFL26* was constructed into a modified pCAMBIA1302 with the kanamycin-resistant gene under the control of the *CaMV 35S* promoter. The *DR5*-*YFP* construct was modified with the replacement of *GFP* by *YFP* from the plasmid of *DR5*-*GFP* (He et al., 2018).

### Genetic transformation, root phenotyping and chemical treatment

*P. tomentosa* was stably transformed using agrobacterium-mediated infiltration of leaf disks as described previously (Jia et al., 2010). PCR genotyping was conducted with the primers of hygromycin/kanamycin-resistant genes for screening positive transgenic plants. Microcutting-propagated transgenic seedlings were transplanted into soil in pots, and cultivated in a greenhouse at 20-25°C with the light of 10 000 lux under a 16 h: 8 h, day: night cycle. For stem-derived AR phenotyping, shoot segments of 3-4 cm with two or three young leaves were cut from sterilized WT and transgenic seedlings and cultivated on sterile WPM (woody plant medium) without any phytohormones (Jia et al., 2010) at 25°C with 16 h of light of 5,000 lx and 8 h of dark. For phenotyping ARs derived from leaf explants, leaves similar in size were cut from 4-week-old sterilized seedlings and incubated on WPM for rooting at 25°C with 16 h of light of 5,000 lx and 8 h of dark. For examining AR formation under hydroponic condition, the upper part of 2-month-old WT and *miR476a*-OE plants with 5-6 leaves was cut off and transferred to Hoagland solution for rooting.

### Crosssectioning and GUS staining

Stem bases from shoot explants were sectioned with a razor blade, and then stained with 0.05% (w/v) toluidine blue for 5 min. The cross-sections were observed and captured using a light microscope (Zeiss). For GUS staining, cross-sections were fixed in acetone for 1 h at 20°C, and then washed twice in double distilled H_2_O. The cross-sections were soaked in GUS staining solution (0.5 M Tris, pH 7.0, 10% Triton X-100 with 1mM X-Gluc (5-bromo-4- chloro-3-indolyl-D-glucuronide)) for 15 min at 37°C in the dark. After reaction, Chl was removed by use of 75% ethanol three times at 65°C. The Chl-free stained stems were observed under an Olympus 566 SZX16 microscope (Tokyo, Japan) and documented using an Olympus DP73 camera.

### RNA in situ hybridization

For preparing probes, 223-bp gene-specific cDNA fragments were amplified for *PtoRFL26* using the primer pair *PtoRFL26*-in situ-fw/rv (Table S1). The probes were labeled using a DIG RNA Labeling Kit (Roche). Section pretreatment, hybridization and immunological detection were performed as described (Sang et al., 2012).

### qRT-PCR

Total RNA was extracted from tissues of 2-month-old *P. tomentosa* plants using a Plant RNeasy Mini Kit (Qiagen). The cDNA was synthesized using the PrimeScript RT reagent Kit with gDNA Eraser (Takara, Dalian, China). Quantitative real time polymerase chain reaction (qRT-PCR) was performed using SYBR Premix ExTaq (Takara, Japan) in a TP800 Real-Time PCR machine (Takara, Japan). The poplar *Ubiquitin* gene was used as the reference gene as an internal standard. The primers used for qRT-PCR are listed in Table S1.

### Degradome sequencing

Degradome libraries were constructed using RNA isolated from shoot cuttings of WT and *miR476a*-OE lines, respectively, for rooting at 3 DAC as previously described (Ma et al., 2010). Briefly, 200ng polyA-enriched RNAs were annealed with biotinylated random primers, and then those with monophosphate at 5’ end ligated to 5’ adaptors, followed by reverse transcription and PCR amplification. The libraries were single-end sequenced using an Illumina HiSeq2500 at Biomarker Technologies (Beijing, China) following the manufacturer’s recommended protocol. Approximately 20 M single-end tags of 50 bp in length were produced for each library.

For data analyses, the tags with low quality and matched to other noncoding RNA such as rRNA, tRNA, and small nuclear RNA were removed. Subsequently, remained tags were aligned to *P. trichocarpa* v3.0 genome, allowing no more than one mismatch. Targets were identified using CleaveLand (version 4.3). The tags that could be mapped to PPR genes of *P. trichocarpa* were aligned to the 11 cloned *RFL* genes of *P. tomentosa* including *PtoRFL1*, *16*, *17*, *18*, *19*, *26*, *27*, *30*, *32*, *33*, and *34*.

### Quantification of ATP content, NAHD/NAD^+^, and H_2_O_2_

Shoot cuttings at 3 DAC during AR formation were sampled for quantifying ATP content, NADH/NAD^+^ and H_2_O_2_. Quantification of ATP and H_2_O_2_ content was conducted using ATP Assay Kit (S0026) and Hydrogen Peroxide Assay Kit (S0038) both produced by Beyotime Biotechnology (Shanghai, China). NADH/NAD^+^ was assayed using NAD(H) Assay Kit (Nanjing Jiancheng Bioengineering Institute, Nanjing, China). The experiments were performed according to the manufacturer’s manuals.

### RNA sequencing

Total RNA was isolated from 5 DAC shoot cuttings of WT and *miR476a*-OE plants using RNeasy Plant Mini Kit (QIAGEN). Messenger RNA was purified and used for Illumina sequencing according to the guide of the Illumina mRNA sequencing sample preparation. Reads were mapped to *Populus trichocarpa* v3.0 genome using Bowtie2 and HISAT allowing up to two mismatches and only uniquely mapped reads were retained. Expression level of loci was determined by fragment per kilobase per million mapped reads (FPKM). Differentially expressed genes (DGEs) were defined as FPKM change in WT and *miR476a*-OE plants of more than 2 folds and FDR < 0.01. The DGEs were then conducted for functional classification using the website of agriGO (http://bioinfo.cau.edu.cn/agriG).

### Gene accessions

GeneBank accession numbers of the *P. tomentosa* genes are: *PtoRFL1* (MN242831), *PtoRFL16* (MN242830), *PtoRFL17* (MN242832), *PtoRFL18* (MN242833), *PtoRFL19* (MN242834), *PtoRFL26* (MN242835), *PtoRFL27* (MN242836), *PtoRFL30* (MN242837), *PtoRFL32* (MN242838), *PtoRFL33* (MN242839), and *PtoRFL34* (MN242840).

## Acknowledgements

We thank Drs Mi Zhang and Jianyan Zeng (Southwest University, China) for auxin content assays. This work was supported by the National Natural Science Foundation of China (31870657, 31800505 and 31870175) and the Fundamental Research Funds for the Central Universities (XDJK2018AA005). We also thank the financial support from the Fundamental Research Funds for the Central Universities of China grant 2572018CL01 and Heilongjiang Touyan Innovation Team Program.

## Author contributions

K.L., C.X. and H.X. designed the work; C.X., Y.T., X.F., L., H.X., H.S., X.W., J.H., and D.F. performed experiments and data analyses; C.X. drafted the manuscript; K.L. and V.C. revised the manuscript. All authors approved the final version of the manuscript for publication.

## Supplemental information

Figure S1. Sequence alignment and secondary structure prediction of poplar *miR476*s.

Figure S2. Additional root phenotypes of *miR476a*-overexpressing transgenic lines.

Figure S3. Generation of STTM-based *miR476a* knockdown transgenic poplar lines.

Figure S4. Phylogenetic analysis of the *RFL*-*PPR* subfamily from *P. trichocarpa*.

Figure S5. Predicted binding sites of *miR476a* harbored in poplar *RFL* genes.

Figure S6. Degradome sequencing and expression of non-*miR476a*-targeted *RFL*s.

Figure S7. Expression of the *miR476a*-targeted *RFL* genes under wounding treatment.

Figure S8. Domain analysis and mitochondrial localization prediction of the PPR proteins encoded by the *miR476a*-targeted *RFL* genes.

Figure S9. RNA in situ hybridization of *RFL26* at 1 and 6 DAC during AR formation in poplar.

Figure S10. Validation of mitochondrial localization and *miR476a*-directed degradation of RFL proteins.

Figure S11. Generation and AR phenotypes of *RFL26m*-overexpressing poplar lines.

Figure S12. Expression levels of mitochondrial-encoded genes in *miR476a*-OE (A) and *RFL26m*-OE (B) lines.

Figure S13. ATP content assays and mitochondrial perturbation during AR formation in Arabidopsis.

Figure S14. Comparative transcriptomic analysis between WT and *miR476a*-OE lines.

Dataset S1. Sequences of degraded fragments identified by degradome sequencing.

Dataset S2. Differentially expressed genes in the RNAseq-based transcriptomic analysis of *miR476a*-OE vs WT.

Table S1. Sequences of oligonucleotide primers and probes used in this study.

